# Dual origins of measured phase-amplitude coupling reveal distinct neural mechanisms underlying episodic memory in the human cortex

**DOI:** 10.1101/084194

**Authors:** Alex P. Vaz, Robert B. Yaffe, John H. Wittig, Sara K. Inati, Kareem A. Zaghloul

**Author notes:** **Correspondence should be addressed to**: Kareem A. Zaghloul Surgical Neurology Branch, NINDS, National Institutes of Health Building 10, Room 3D20 10 Center Drive Bethesda, MD 20892-1414.

## Abstract

Phase-amplitude coupling (PAC) is hypothesized to coordinate neural activity, but its role in successful memory formation in the human cortex is unknown. Measures of PAC are difficult to interpret, however. Both increases and decreases in PAC have been linked to memory encoding, and PAC may arise due to different neural mechanisms. Here, we use a waveform analysis to examine PAC in the human cortex as participants with intracranial electrodes performed a paired associates memory task. We found that successful memory formation exhibited significant decreases in left temporal lobe and prefrontal cortical PAC, and these two regions exhibited changes in PAC within different frequency bands. Two underlying neural mechanisms, nested oscillations and sharp waveforms, were responsible for the changes in these regions. Our data therefore suggest that decreases in measured cortical PAC during episodic memory reflect two distinct underlying mechanisms that are anatomically segregated in the human brain.

## Introduction

Interactions between oscillations in the brain remain poorly understood, though they are hypothesized to temporally coordinate information between local neuronal populations (J. Lisman & Idiart, 1995; J. Lisman, 2005; Jensen & Colgin, 2007; Canolty & Knight, 2010; J. E. Lisman & Jensen, 2013). One such interaction, phase-amplitude coupling (PAC), occurs when the phase of a low frequency oscillation modulates the amplitude of a high frequency oscillation (Canolty et al., 2006; Canolty & Knight, 2010). In the context of memory, this may provide a mechanism for embedding, and retrieving, individual memory representations within a broader context (Hasselmo & Eichenbaum, 2005; Buzs´aki, 2005; Canolty & Knight, 2010). Indeed, evidence that PAC may play a role in memory formation has emerged in studies of the hippocampus in both animals and humans (Tort, Komorowski, Manns, Kopell, & Eichenbaum, 2009; Axmacher et al., 2010; J. E. Lisman & Jensen, 2013; Lega, Burke, Jacobs, & Kahana, 2014; Heusser, Poeppel, Ezzyat, & Davachi, 2016).

Despite empiric support for the role of PAC in memory, however, successful memory encoding has also been linked with decreases, rather than just increases, in PAC (Lega et al., 2014; Axmacher et al., 2010; Leszczynski, Fell, & Axmacher, 2015). This raises the question as to whether, in some cases, PAC limits effective neural processing. This may be the case in the cortex, where PAC has been observed in pathologic conditions such as Parkinson’s disease (de Hemptinne et al., 2013, 2015). Hence, although PAC is ubiquitous throughout the cortex (Canolty et al., 2006; He, Zempel, Snyder, & Raichle, 2010; Canolty & Knight, 2010), it is unclear whether cortical PAC is beneficial for memory encoding. One possibility is that increases in cortical PAC may improve memory encoding, lending support to the hypothesis that cortical PAC coordinates information just as in the hippocampus. Conversely, if PAC actually limits information processing in the cortex, then successful memory formation should be accompanied by decreases in PAC.

We examine this question here using intracranial EEG (iEEG) in participants with subdural electrodes placed for seizure monitoring as they engaged in a paired associates verbal memory task. Importantly, properly interpreting these data must account for the fact that measured PAC may arise due to two different underlying neural mechanisms - true interactions between low and high frequency oscillations that may help coordinate local neural populations (Jensen & Colgin, 2007; J. E. Lisman & Jensen, 2013), or repeated sharp or non-sinusoidal deflections in the iEEG signal resulting in broadband increases in power that may be related to general neural activation (Makeig et al., 2002; Manning, Jacobs, Fried, & Kahana, 2009; Miller et al., 2007; Burke, Ramayya, & Kahana, 2015; Lozano-Soldevilla, ter Huurne, & Oostenveld, 2016). As such, we used a waveform analysis to investigate the electrophysiological contributions to any changes in PAC. We were motivated to understand whether any changes in cortical PAC related to successful memory encoding could be attributed to these distinct neural mechanisms.

## Methods

### Participants

33 participants with medication-resistant epilepsy underwent a surgical procedure in which platinum recording contacts were implanted subdurally on the cortical surface as well as deep within the brain parenchyma. In each case, the clinical team determined the placement of the contacts to localize epileptogenic regions. The Institutional Review Board (IRB) approved the research protocol, and informed consent was obtained from the participants and their guardians. These data were initially collected and analyzed for changes in spectral power in separate studies (Yaffe et al., 2014; Greenberg, Burke, Haque, Kahana, & Zaghloul, 2015).

### Paired Associates Task

Each patient participated in a paired associates task (Figure 1a). Participants were asked to study a list of word pairs and then later cued with one word from each of the pairs, selected at random. Participants were instructed to vocalize each cue word’s partner from the corresponding word pair. Lists were composed of four pairs of common nouns, chosen at random and without replacement from a pool of high-frequency nouns. Words were presented sequentially and appeared in capital letters at the center of the screen. Word pairs were separated from their corresponding recall cue by a minimum lag of two study or test items. During the study period (encoding), each word pair was preceded by an orientation stimulus (a row of capital X’s) that appeared on the screen for 300 ms followed by a blank interstimulus interval (ISI) of 750 ms with a jitter of 75 ms. Word pairs were then presented on the screen for 2500 ms followed by a blank ISI of 1500 ms with a jitter of 75 ms. During the test period (retrieval), one randomly chosen word from each study pair was shown, and the participant was asked to recall the other word from the pair by vocalizing a response into a microphone. Each cue word was preceded by an orientation stimulus (a row of question marks) that appeared on the screen for 300 ms followed by a blank ISI of 750 ms with a 75 ms jitter. Cue words were then presented on the screen for 3000 ms followed by a blank ISI of 4500 ms. Participants could vocalize their response any time during the recall period after cue presentation. Vocalizations were digitally recorded and then manually scored for analysis. Responses were designated as correct, as intrusions, or as passes when no vocalization was made or when the participant vocalized the word ‘pass’. Intrusion and pass trials were designated as incorrect trials. A single experimental session contained up to 25 lists. For analysis, we designated a single trial as the encoding period for a study word pair and the retrieval period during testing of its corresponding cue.

**Figure 1.**
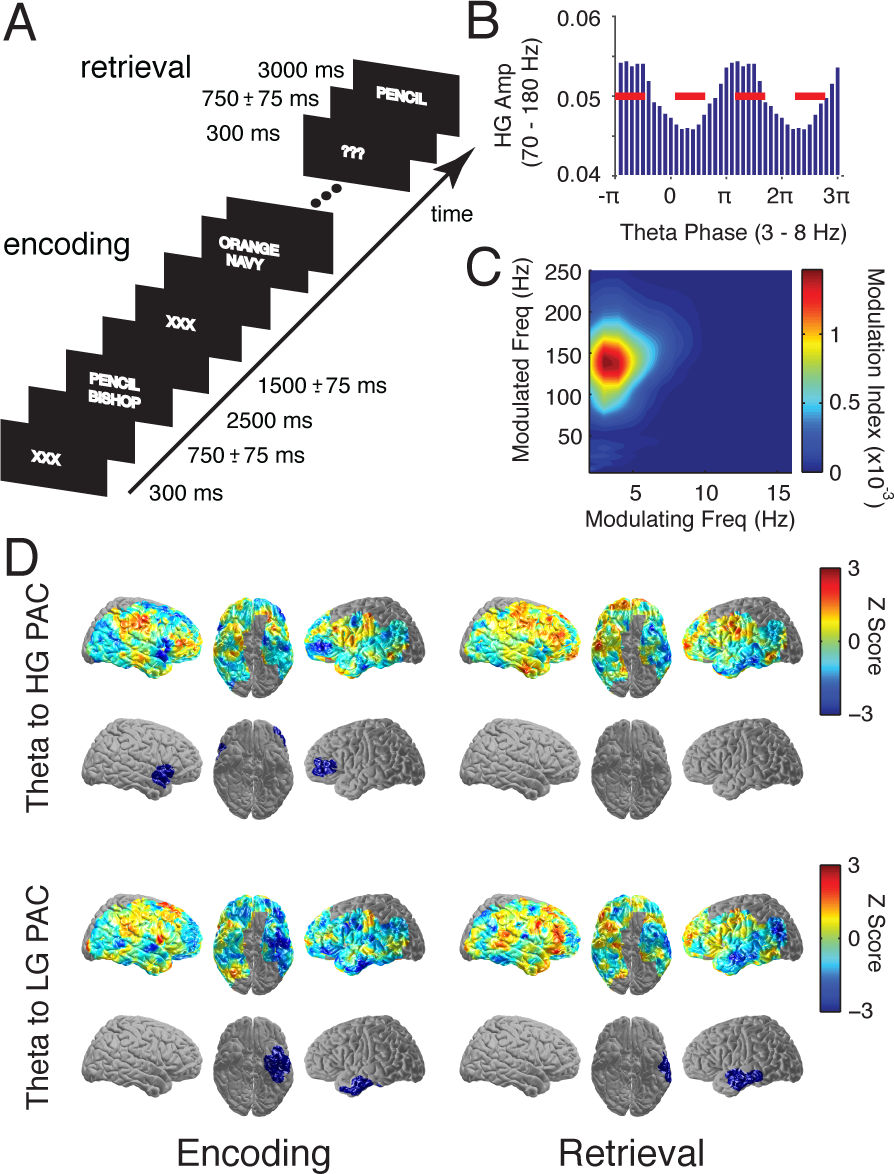
Methods and PAC analysis (**A**) Paired associates episodic memory task. In each trial, a pair of words is presented on the screen during encoding. Subsequently during retrieval, one word from each pair is randomly chosen and presented, and participants are instructed to vocalize the associated word. Timing of word and cue presentation is shown in the inset. (**B**) Theta to HG (70-180 Hz) phase-amplitude plot. Average normalized HG amplitude for this electrode is plotted for 20 theta (3-8 Hz) phase bins repeated for two cycles. Dotted red line indicates HG amplitude where no comodulation would be measured. (**C**) Comodulogram. Each frequency pair of phase providing frequencies every 1 Hz from 2 Hz to 16 Hz versus amplitude providing frequencies every 5 Hz from 5 Hz to 250 Hz is plotted where colors correspond to the respective modulation indices (MI) of each pair. (**D**) Whole brain analysis across participants for theta to LG and HG PAC for correct versus incorrect trials. Colors in the first and third rows correspond to the average *z*-score differences between correct and incorrect trials in each ROI during encoding and retrieval. ROIs exhibiting significant differences, corrected for multiple comparisons, are shown in the second and fourth rows. The blue color of the significant regions in these rows indicates PAC was significantly less for correct trials compared to incorrect trials.

### Intracranial EEG (iEEG) Recordings

Depending on the amplifier and the discretion of the clinical team, intracranial EEG (iEEG) signals were sampled at 1000 or 2000 Hz. Signals were referenced to a common contact placed subcutaneously, on the scalp, or on the mastoid process. All recorded traces were resampled at 1000 Hz, and a fourth order 2 Hz stopband butterworth notch filter was applied at 60 Hz to eliminate electrical line noise. The testing laptop sent +/-5 V digital pulses via an optical isolator into a pair of open lines on the clinical recording system to synchronize the electrophysiological recordings with behavioral events.

We collected electrophysiological data from a total of 2750 subdural and depth recording contacts (PMT Corporation, Chanhassen, MN; AdTech, Racine, WI). Subdural contacts were arranged in both grid and strip configurations with an inter-contact spacing of 10 mm. Hippocampal depth electrodes (6-8 linearly arranged contacts) were placed in four patients. Contact localization was accomplished by co-registering the post-op CTs with the post-op MRIs using both FSL Brain Extraction Tool (BET) and FLIRT software packages and mapped to both MNI and Talairach space using an indirect stereotactic technique and OsiriX Imaging Software DICOM viewer package. The resulting contact locations were subsequently projected to the cortical surface of a Montreal Neurological Institute N27 standard brain (Dykstra et al., 2011). Pre-operative MRIs were used when post-operative MR images were not available.

We analyzed iEEG data using bipolar referencing to reduce volume conduction and confounding interactions between adjacent electrodes (Nunez & Srinivasan, 2006). Bipolar referencing is routinely used for subdural electrode recordings, and has been noted be superior to the average reference montage in reducing muscular artifacts in iEEG (Kovach et al., 2011). We defined the bipolar montage in our data-set based on the geometry of iEEG electrode arrangements. For every grid, strip, and depth probe, we isolated all pairs of contacts that were positioned immediately adjacent to one another; bipolar signals were then found by finding the difference in the signal between each pair of immediately adjacent contacts. The resulting bipolar signals were treated as new virtual electrodes (henceforth referred to as electrodes throughout the text), originating from the midpoint between each contact pair. All subsequent analyses were performed using these derived bipolar signals. Importantly, we excluded all electrodes exhibiting ictal or inter-ictal activity at the seizure focus and at sites of generalization as identified by a team of trained epileptologists in order to avoid confounding any memory effects with concurrent seizure activity. In total, our dataset consisted of 2,292 electrodes (1,048 left hemispheric, 1,244 right hemispheric). Additionally, we excluded any trials displaying excessive variance or kurtosis (defined as greater 2.3 times the interquartile range away from the third quartile) in order to provide the most conservative assessment of normal human electrophysiology in the context of noise and transient epileptiform activity.

### Spectral Power and Phase

We quantified spectral power and phase by convolving the bipolar iEEG signals with complex valued Morlet wavelets (wavelet number 6) (Addison, 2002). To quantify phase-amplitude coupling (PAC) and to generate corresponding comodulograms, we calculated spectral power and phase using 15 linearly spaced wavelets between 2 and 16 Hz for the modulating frequencies and 25 linearly spaced wavelets between 5 and 250 Hz for the modulated frequencies. The encoding period was 0 to 4500ms from the onset of pair presentation, and the retrieval period was from the onset of cue presentation to the time of vocalization. We convolved the above wavelets with the iEEG data from each of these periods in order to generate continuous measures of instantaneous amplitude and phase. During pass trials where no vocalization was present, we assigned a response time by randomly drawing from the distribution of correct reaction times. One participant was instructed to vocalize her response only after the cue word disappeared from the screen. In all trials, we included a 1000 ms buffer on both sides of the clipped data.

To examine PAC between frequency bands, we also quantified spectral power and phase for each frequency band by first bandpass filtering the iEEG signal into four predefined frequency bands using a second order Butterworth filter: theta (3-8 Hz), alpha (8-12 Hz), low gamma (30-58 Hz), and high gamma (70-180 Hz). We then calculated a continuous measure of amplitude and phase for each frequency band using a Hilbert transform. While the delta band (1-3 Hz) may be important in specific memory contexts (Haque, Wittig Jr., Damera, Inati, & Zaghloul, 2015), we did not analyze delta band PAC due to the reported intrinsic bias of PAC towards lower frequencies (Aru et al., 2015). Non-stationarities of the iEEG signal occupy less phase bins than those of higher frequency modulating bands. Hence it follows that delta PAC measurements are more vulnerable to nonspecific high frequency fluctuations in the iEEG signal (Aru et al., 2015; Jones, 2016), and we report theta and alpha band effects instead.

### Measurement of Phase-Amplitude Coupling

We used the continuous measures of phase and amplitude to compute phase-amplitude coupling (PAC) between every modulating low frequency and every modulated high frequency. During every temporal epoch (i.e. encoding or retrieval), we divided the continuous time phases of the modulating frequency into 20 evenly spaced phase bins. For every phase of the modulating low frequency signal, we found the corresponding amplitude of the modulated high frequency signal and assigned that amplitude to the corresponding phase bin. The presence of PAC results in a non-uniform distribution of modulated amplitudes across modulating phase bins. We quantified this by calculating a modulation index (MI) which measures the difference in entropy between the calculated phase-amplitude distribution and a uniform distribution using a normalized Kullback-Leibler distance (Tort, Komorowski, Eichenbaum, & Kopell, 2010):

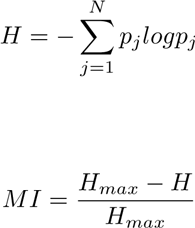

where *j* is the index of each phase bin, *N* is the total number of phase bins, and *H* is the entropy of the distribution *p_j_. H*_*max*_ is the entropy of a uniform phase-amplitude distribution and is defined as *logN*. We constructed comodulograms by calculating the MI between every pair of low modulating frequencies and high modulated frequencies. For every calculated MI, we defined the preferred phase as the phase bin with the highest average modulated amplitude.

### Anatomic Localization and Topographic Plots

With iEEG, the precise placement of electrodes is different for each participant, which limits our ability to examine spatially resolved effects across subjects. We overcome this limitation by spatially smoothing electrode effects using 760 spatial regions of interest (ROIs) that were evenly spaced every 9.98 ± 0.02 mm on the cortical surface of a Montreal Neurological Institute N27 standard brain. For each participant, we averaged either the z-scored MI or waveform amplitude from all electrodes that were within 12.5 mm from the center of a given ROI. We averaged statistics within each individual so that a single region was not overly represented by a participant who happened to have many electrodes in that region. Most electrodes contributed to more than one ROI, and most ROIs included either zero or more than one electrode per subject. When performing whole brain analyses across participants, only ROIs that contained electrodes from five or more participants were evaluated.

We visualized ROI-based results by projecting the values computed at each ROI (see Statistical Analyses) onto a 3D rendered image of the standard brain. For each vertex of the standard brain, a weighted average of nearby ROI values was calculated using a 3D gaussian kernel (*σ* = 4.2 mm; center weight 1; zero weight beyond 12.5 mm). With this method, a vertex directly adjacent a particular ROI will largely represent the value of that ROI, with a small contribution from neighboring ROIs (which are spaced approximately 10 mm apart). On the other hand, a vertex halfway between two ROIs will represent an attenuated mean of those two ROIs. For whole-brain plots demonstrating significant differences between trial types, all colored regions, independent of color intensity, indicate two-tailed significance at the *p* < 0.05 level as determined by the non-parametric cluster-based correction for multiple comparisons described below.

### Statistical Analyses

For every participant, we calculated the MI across all trials for every electrode. We did this separately for correct and incorrect trials to generate a difference in MI for every pair of modulating and modulated frequency bands during each temporal period (encoding or retrieval). To assess for significant differences between conditions, we first permuted the trial labels 1000 times and generated an empiric distribution of MI differences in each electrode for each participant. The true difference was then assigned a *z*-score based on the normal inverse cumulative distribution function (*µ* = 0, *σ* = 1) of its rank among the permuted values. Hence, for each electrode, we generate a measure of statistical significance (|*z*| > 1.96, *p* < .05) when comparing PAC between correct and incorrect trials. In addition to providing a measure of significance, this normalization procedure allows for the comparison of MI value changes that can vary several orders of magnitude across electrodes. We then spatially smoothed each electrode’s *z*-scored MI difference by converting to ROIs as described above. For every ROI in each participant, we were therefore left with an average *z*-scored difference in MI during each temporal epoch and for every pair of frequency bands.

We performed a random effects statistical analysis at each ROI across participants. Our null hypothesis was that across participants, the brain region represented by each ROI showed no difference in PAC during correct versus incorrect trials. We tested this hypothesis using a non-parametric clustering-based procedure to assess for statistical differences across participants (Maris & Oostenveld, 2007). This procedure identifies contiguous spatial regions exhibiting significant differences between trial types while avoiding a priori assumptions about particular spatial regions and correcting for multiple comparisons. For each analyzed ROI, a minimum of five participants contributed across-trial *z*-scored MI difference values. In each ROI, we computed the true mean difference across participants between correct and incorrect trials using these values. We then randomly permuted the participant-specific averages (correct vs incorrect), which in practice translates to randomly reversing the sign of the *z*-scored MI difference, within each participant and recomputed the mean difference across participants. For *n* participants, this results in an empiric distribution of 2_*n*_ possible mean differences that are all equally probable under the null hypothesis. We generated the empiric distribution from a minimum of 32 (number of participants = 5) and a maximum of 1000 permutations for every ROI. We compared the true mean difference in each ROI to this empiric distribution to generate an across-participant *z*-score and *p*-value for each ROI. This *p*-value represents the likelihood that the true mean difference at an individual ROI represents a departure from the null hypothesis. However, this *p*-value for each individual ROI does not take into account the multiple comparisons that are made in space (across ROIs), and therefore is not reported in the text.

To correct for multiple comparisons across ROIs, we separately identified positive and negative clusters containing ROIs that were adjacent in space that exhibited a significant difference between trial types using the across-participant permutation procedure described above (where in each ROI, *p* < .05). For each cluster of significant ROIs identified in the true and permuted cases, we defined a cluster statistic as the sum of each of the *z*-scores for all ROIs within that spatial cluster. We retained the maximum cluster statistic during each permutation to create a distribution of cluster statistics. We assigned *p*-values to each identified cluster of the true data by comparing its cluster statistic to the distribution of maximum cluster statistics from the permuted cases. Clusters were determined to be significant if their *p*-value calculated in this manner was less than .05.

### Generation of Average Waveforms

To identify a characteristic waveform in the iEEG signal in a particular electrode, we first identified the low frequency modulating and high frequency modulated frequency band associated with the highest MI and its associated preferred phase bin. We examined the bandpass filtered and Hilbert transformed iEEG signal and identified all instances of this preferred phase. Because the preferred phase is in fact a phase bin that extends over a discrete temporal window, we identified the precise time of maximum high gamma amplitude within each temporal window and designated that point as the time of the preferred phase.

Then at every instance of the preferred phase in the modulating frequency, we extracted 500 ms of the continuous iEEG signal centered at the precise time of maximum high gamma amplitude. We averaged these extracted waveforms across all instances of preferred phase across all trials to generate an average waveform for each electrode during each condition. In this manner, average waveforms with larger amplitudes reflect either a larger or more consistent occurrence of a characteristic waveform during the preferred phase of each cycle of the modulating signal.

We used an automated algorithm to categorize the type of waveform found in each electrode. We categorized a waveform as a nested oscillation if at least three local maxima fell within 3 cycles of the slowest possible high frequency oscillation of the modulated frequency band. In practice, this meant finding 3 local maxima around the preferred phase within 100 ms for low gamma electrodes or within 45 ms for high gamma electrodes, which have the slowest possible frequencies of 30 Hz and 70 Hz respectively. Electrodes that did not satisfy this criteria were categorized as sharp waveforms.

To confirm that nested oscillation electrodes were designated based on genuine oscillations as opposed to random high frequency fluctuations in the signal, we generated surrogate waveforms by selecting a random preferred phase for each trial and then implementing the same triggered average. In this fashion, we generated the surrogate preferred phase waveforms with the same temporal autocorrelations as the true preferred phase waveforms. We assigned the sizes of the local maxima in all waveforms as the distance from the peak to the nearest local minima, and we performed a pairwise t-test comparing the mean local maxima size of each trial to its surrogate for each electrode. We found that the high frequency peaks of the nested oscillations were significantly (*p* < .05) larger than those of their surrogate counterparts in 86.39 ± 1.73 % of electrodes across all participants. We designated nested oscillations electrodes that had high frequency components that were not significantly different from their surrogates as indicative of stochastic volatility of the signal, and did not include them as nested oscillation electrodes in subsequent analyses.

### Calculation of Sample Entropy

Due to the apparent cyclostationarity of sharp waveforms as opposed to nested oscillations, we investigated if sample entropy, a measure of signal complexity, was as anatomically tractable as waveform type. For a given template sequence of length *m+1*, where *m* is known as the embedding dimension, sample entropy can be understood as the cumulative probability that one can predict *m+1* length self-similar template sequences based on *m* length self-similar sequences. Sample entropy therefore measures signal complexity by quantifying the occurrence of self-similar sequences in the signal. Intuitively then, less predictable signals (i.e. less cyclostationary) are expected to have higher values of sample entropy.

For an embedding dimension *m*, a tolerance *r*, and a time-step *τ*, the formal equations for the calculation of sample entropy are as follows (Sokunbi et al., 2013):

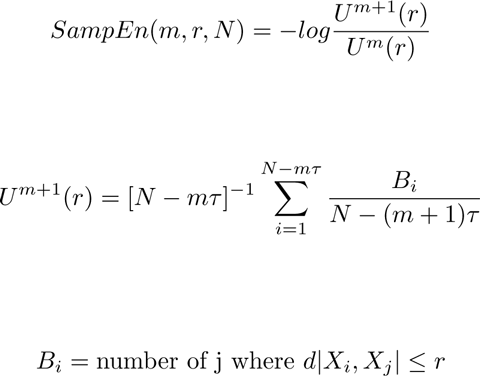

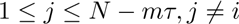

*X*^*i*^ and *X*^*j*^ are pattern vectors whose components are time delayed versions of the original signal *X*(*t*) with time-step τ. An embedding dimension *m* of 2, a tolerance *r* of 0.2 * *std*(*X*(*t*)), and a constant time-step *τ* of 1 ms were used in all analyses. Of note, the number of 3 element self-similar template sequences is necessarily less than or equal to the number of 2 element self-similar template sequences, implying that sample entropy is bounded between 0 and 1.

## Results

33 participants (18 males; age 34.2 ± 2.3 years (mean ± SEM)) with medically refractory epilepsy who underwent surgery for placement of intracranial electrodes for seizure monitoring participated in a verbal paired associates task (Figure 1a). Participants responded on each trial with the correct word, an incorrect word (intrusion), or with no word. Responses were designated as passes when either no response was given or the word ‘pass’ was vocalized. We considered intrusion and pass trials as incorrect trials for all analyses. Participants studied 224 ± 23 word pairs and successfully recalled 38.3 ± 3.6 % of paired words with a mean response time of 1,812 ± 75 ms. Participants vocalized intrusions on 14.6 ± 1.9 % of trials with a mean reaction time of 2,722 ± 112 ms. For the remaining study word pairs, participants either made no response to the cue word, or vocalized the word ‘pass’ with a mean response time of 3,385 ± 200 ms. A one-way ANOVA showed that study-test lag had no significant effect on recall probability (*F* (4, 160) = .97, *p* = .43).

We analyzed intracranial EEG (iEEG) oscillatory activity to investigate interactions between low frequency and high frequency signals. We used a measure of phase-amplitude coupling (PAC) to quantify the extent to which the phase of a low frequency oscillation is coupled to the amplitude of a higher frequency (Tort et al., 2008, 2009, 2010; Lega et al., 2014). In many electrodes across participants, we found that high frequency amplitude was locked to the phase of a low frequency oscillation. We define the phase at which the peak of the high frequency amplitude occurs as the preferred phase (Figure 1b). For every electrode, we constructed a phase-amplitude comodulogram by calculating PAC for every pair of low and high frequencies (Figure 1c). We separately constructed these comodulograms for correct and incorrect trials during the encoding and retrieval periods to assess any differences in PAC between conditions.

Based on the known interactions between theta and gamma frequencies in the hippocampus during memory formation (Axmacher et al., 2010; Lega et al., 2014), we examined whether similar interactions between both theta and low gamma (30-58 Hz) and theta and high gamma (70-180 Hz) existed throughout cortical regions. We also examined alpha to low and high gamma interactions given the role that alpha to high gamma PAC may play during tasks that involve visual processing, attention, and memory (Palva, Palva, & Kaila, 2005; Cohen et al., 2009; Canolty & Knight, 2010; Voytek et al., 2010; Fell & Axmacher, 2011). Across participants, we found that 37.86 ± 1.79 % of all electrodes demonstrated significant differences in PAC between correct and incorrect trials in any of these frequency band pairs in either encoding or retrieval (*p* < .05, permutation procedure). 11.86 ± 1.06 and 13.30 ± 1.73 % of electrodes demonstrated significant differences in theta to low gamma and theta to high gamma PAC, respectively, between conditions, while 10.81 ± 0.67 and 11.91 ± 1.06 % of electrodes demonstrated significant differences in alpha to low or high gamma PAC, respectively.

We examined the spatial distribution of these significant effects by spatially smoothing electrode data from each participant into regions of interest (ROIs) that spanned the entire brain (see Methods). We identified contiguous clusters of ROIs that exhibited a significant difference in PAC between correct and incorrect trials across participants (*p* < .05, permutation procedure, corrected for multiple comparisons; see Methods). Theta band PAC to both low and high gamma exhibited significant memory related changes in distinct anatomic areas. During encoding, correct trials had significantly decreased theta to high gamma PAC in the left and right inferior frontal gyri, including a small portion of the right superior temporal gyrus (Figure 1d). Conversely, theta to low gamma PAC significantly decreased in the left temporal lobe during both encoding and retrieval. For alpha band PAC, we found significant decreases in alpha to low gamma PAC bilaterally in the medial temporal lobes along the parahippocampal gyri and in the right orbitofrontal cortex during encoding (data not shown). These differences, however, were not present in any brain region for retrieval, and no significant regions were found for alpha to high gamma coupling in either encoding or retrieval.

Measures of PAC may arise due to true interactions between low and high frequency oscillations (Jensen & Colgin, 2007; J. E. Lisman & Jensen, 2013) or due to repeated phase-locked sharp deflections in the recorded trace (Kramer, Tort, & Kopell, 2008; Aru et al., 2015; Jones, 2016; Lozano-Soldevilla et al., 2016). As such, we were interested in understanding whether differences in these underlying neural mechanisms giving rise to PAC could account for the observed differences in these anatomic distributions. We began by examining the relationship between the unfiltered iEEG voltage signal and the observed measure of PAC in all electrodes. Representative examples of the iEEG signal, the filtered low and high frequency signal, the spectrogram, and the corresponding comodulogram for three electrodes are shown in Figure 2. Consistent with the conventional interpretation of PAC (Canolty et al., 2006; Buzs´aki, 2006; J. E. Lisman & Jensen, 2013), we found that PAC may emerge from the interaction between a low frequency signal and a phase-locked high frequency oscillation (Figure 2a). We call this type of iEEG signal a nested oscillation, as multiple cycles of a high frequency oscillation are locked to a low frequency waveform. However, we also observed that apparent PAC could emerge in the presence of a repeated sharp waveform that, although containing high frequency components, often does not exhibit an explicit high frequency oscillation (Figure 2b-c). We call these iEEG signals sharp waveforms. Notably, these waveforms can at times consist of only a single low frequency oscillation with sharp peaks that generate high frequency components (Figure 2b).

**Figure 2.**
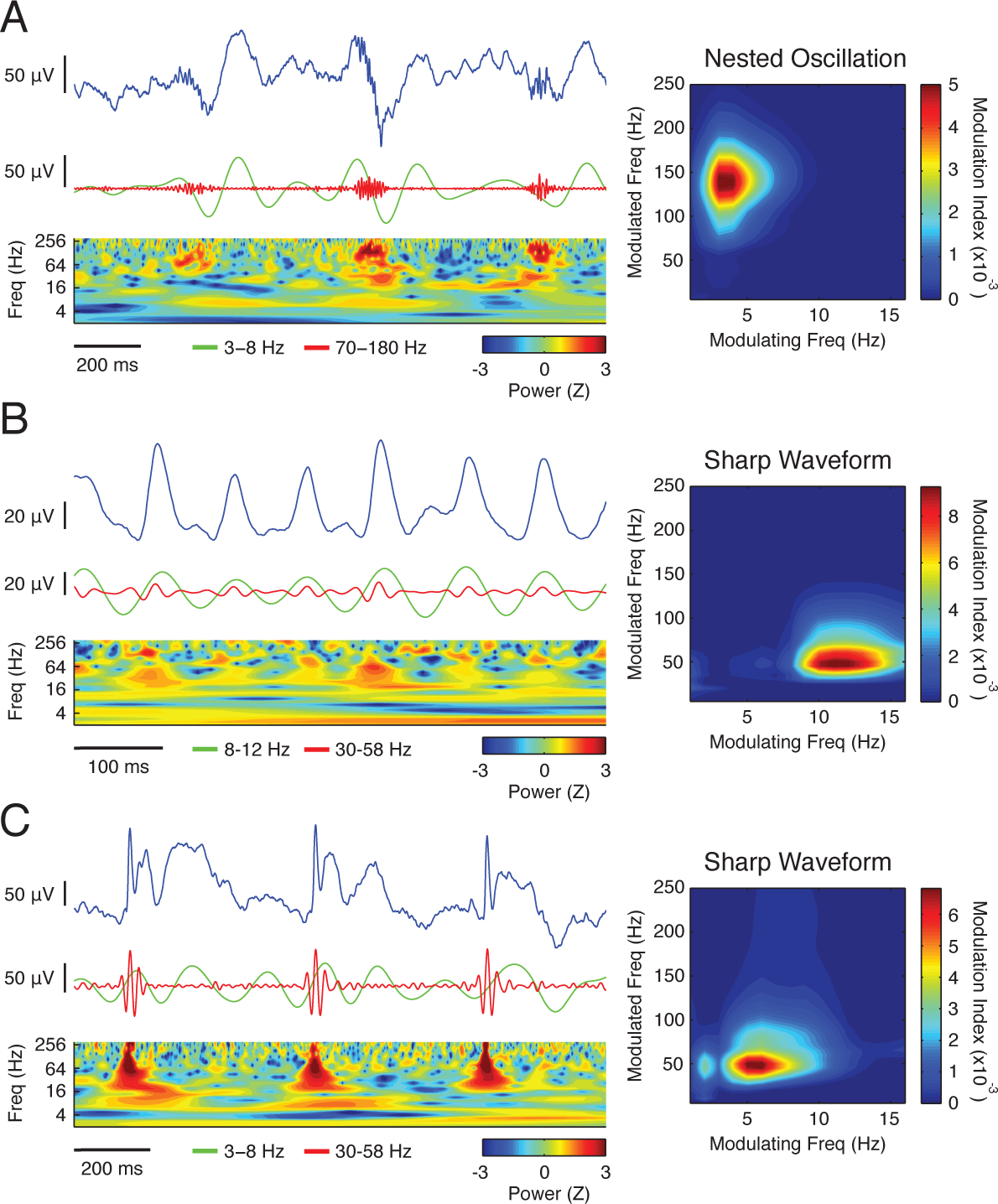
Characteristic waveforms underlie PAC (**A**) Nested oscillation (NO) example of PAC. (*left*) Exemplar iEEG trace is shown in blue while low and high frequency bandpassed components are shown in green and red respectively. (*right*) Corresponding comodulograms are shown for all trials for each electrode. (**B**) Sharp waveform (SW) example of PAC from single low frequency oscillation. (**C**) Generalized SW example of PAC.

Given this relation between a characteristic waveform and measured PAC, we hypothesized that the extent to which these waveforms are repeated in the iEEG trace could track the observed changes in PAC during successful memory encoding and retrieval. For each electrode, we used the modulating frequency band and the preferred phase identified in the comodulogram to determine the characteristic waveform present in the iEEG trace during each condition (Figure 3a; see Methods). The amplitude of the phase-triggered average waveform reflects a combination of both the consistency and the strength of the modulating low frequency signal. This amplitude should not be significantly impacted by the presence or absence of any high frequency oscillations. We found that, in an exemplar electrode, the amplitude of the average waveform is significantly correlated with the extent to which PAC was measured across all trials (*r* = 0.663, *p* < .001; Figure 3b). This relation between the *z*-scored characteristic waveform amplitude and apparent PAC was consistent across all electrodes in this participant 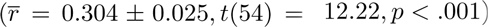, and across all participants 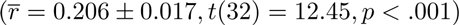 (Figure 3b).

**Figure 3.**
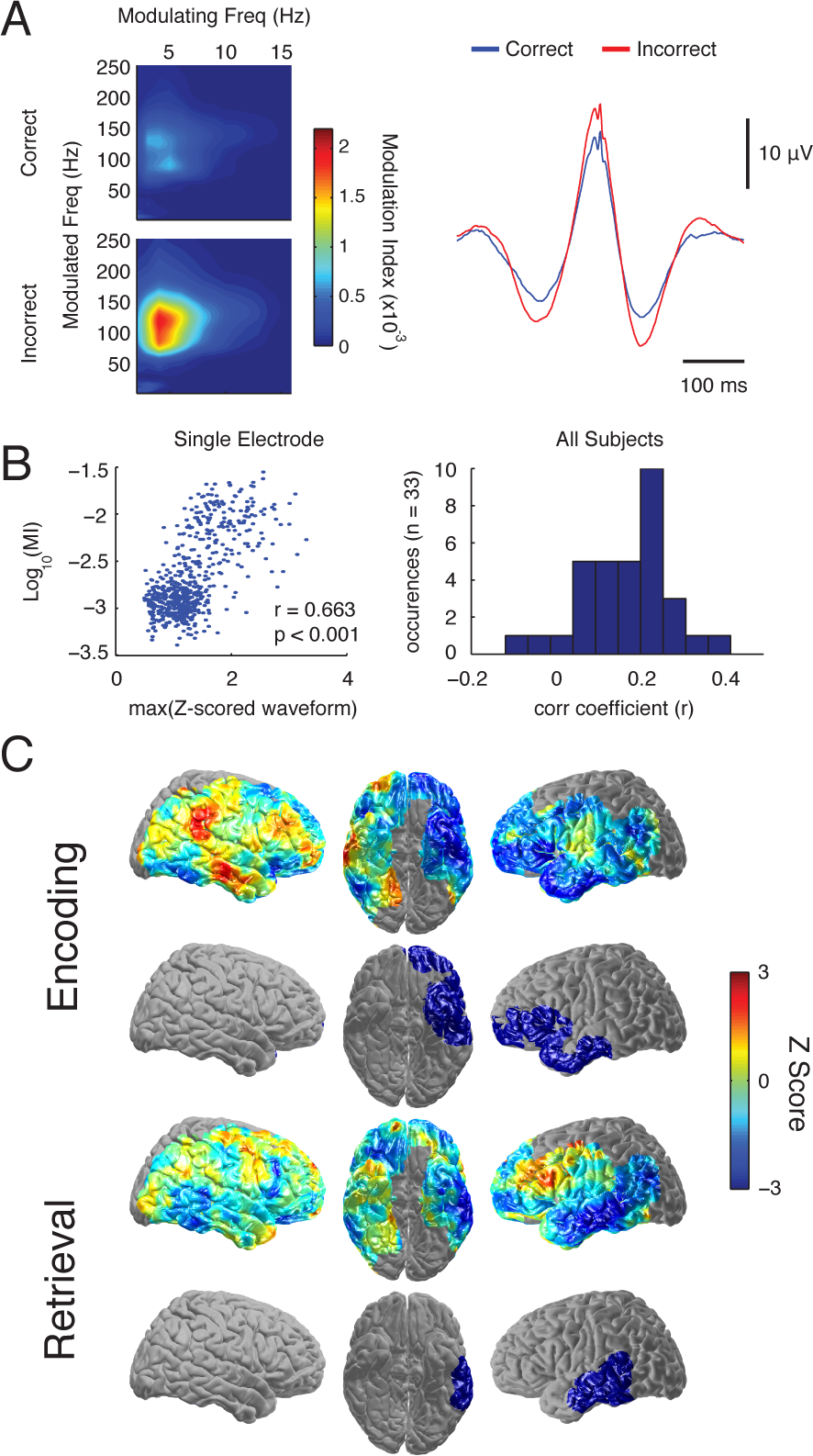
Cortical waveforms exhibit memory related differences (**A**) Comodulograms for correct and incorrect encoding periods with corresponding average waveforms. The average waveforms for correct and incorrect trials are shown on the *right*. (**B**) Correlation of PAC and waveform amplitude. (*left*) Correlation of log-transformed MI vs. amplitude of average *z* scored waveform for all trials of the electrode in (A). (*right*) Average correlation coefficient (*r*) for all participants (n=33). (**C**) Whole brain group analysis for average waveform amplitude for correct compared to incorrect trials. Colors in the first and third rows correspond to the average *z*-score differences between correct and incorrect trials in each ROI during encoding and retrieval. ROIs exhibiting significant differences, corrected for multiple comparisons, are shown in the second and fourth rows. The blue color of the significant regions in these rows indicates waveform amplitude was significantly less for correct trials compared to incorrect trials.

We then examined the memory related differences in these average waveforms between correct and incorrect trials across all electrodes and across all participants. We trial-matched correct and incorrect trials, calculated the difference in peak amplitude between each case for every electrode, and compared the difference across participants. We found contiguous clusters of ROIs that exhibited significant differences in average waveform amplitude between correct and incorrect trials (*p* < .05, permutation procedure, corrected for multiple comparisons; Figure 3c; see Methods). During successful encoding, we found significant decreases in average waveform amplitude in the left inferior frontal gyrus and temporal lobe. During retrieval, however, these differences were only present in the left temporal lobe.

Our data suggest that a large component of the changes in measured PAC observed with memory formation arise due to differences in the consistency and strength of an underlying iEEG waveform. However, the average waveform amplitudes fail to explain how the changes in cortical PAC specific to separate frequency band pairs are differentially distributed across brain regions. We therefore classified each electrode as showing either nested oscillations or sharp waveforms by applying an automated algorithm to the average waveforms extracted from the iEEG trace for each electrode (see Methods). As suggested by previous studies (Kramer et al., 2008; Tort, Scheffer-Teixeira, Souza, Draguhn, & Brankačk, 2013), average waveforms only exhibit high frequency oscillations if the underlying iEEG signal contains an actual high frequency oscillation. Hence, we designated an electrode as exhibiting a nested oscillation if its average waveform contained at least three local maxima around the preferred phase. If three local maxima were not present on the average waveform, we categorized it as exhibiting a sharp waveform. A single subject example of this classification scheme is illustrated in Figure 4.

**Figure 4.**
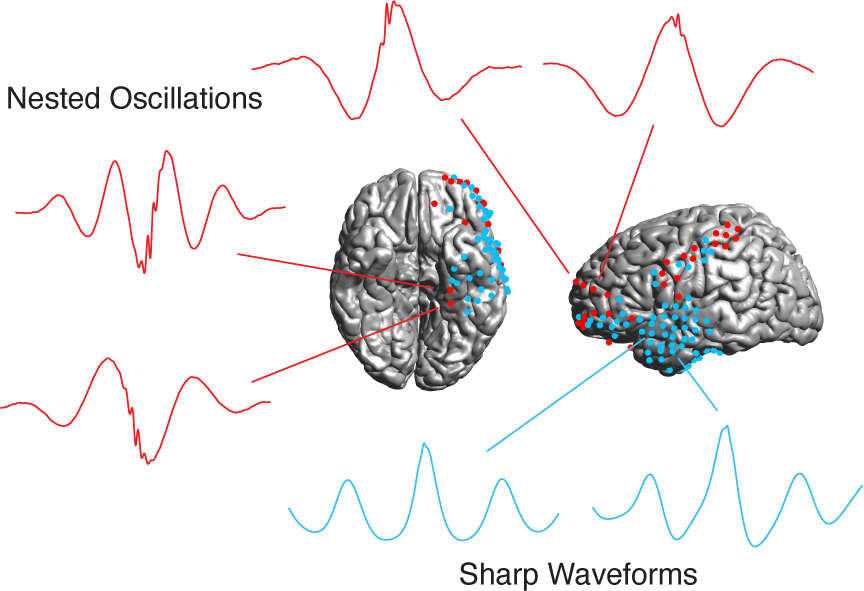
Average waveforms from a single subject. All electrodes for this subject are shown in either red (indicating NO) or blue (indicating SW). Example triggered waveforms are shown from the left parahippocampal gyrus, frontal lobe, and temporal lobe.

We found that approximately a third of all electrodes dominated by a theta modulating frequency were categorized as containing nested oscillations (36.2 ± 3.7% across participants). This was about three times more likely than the appearance of nested oscillations in electrodes dominated by an alpha modulating frequency (11.1 ± 2.7%; *t*(32) = 7.05, *p* < .001). Conversely, slightly over half of all electrodes exhibiting high gamma PAC were categorized as containing nested oscillations (52.9 ± 4.0%). This was also around three times more likely than in electrodes exhibiting low gamma PAC (15.1 ± 2.4%; *t*(32) = 11.00, *p* < .001). Both nested oscillation and sharp waveform electrodes exhibited a non-uniform distribution of phases (nested oscillations, *p* < .001, *z* = 41.04, Rayleigh Test with angle doubling to account for bimodal distribution; sharp waveforms, *p* < .001, *z* = 60.61). The distribution of preferred phases for both categories clustered around both 0 and ±*π*, suggesting that any high frequency components are related to the peaks and troughs of those signals. Of note, however, bipolar referencing may result in preferred phases for adjacent electrodes that are offset by *π*, and so our data cannot determine whether the preferred phases are truly bimodal, or if they only occur at either the peak or trough of the low frequency oscillation.

We next examined the distribution of waveform categories in all electrodes in each spatial ROI (Figure 5a). Electrodes in the left prefrontal cortex and the right occipito-temporal lobe were more likely to be classified as containing nested oscillations, while electrodes in the temporal lobes were more likely to be classified as demonstrating sharp waveforms. Indeed, we found significant differences in the distribution of electrode categories across frontal, temporal, and parietal lobe electrodes (*f*(72) = 11.72, *p* < .001, one-way ANOVA across brain areas for each waveform category). In post-hoc tests, we found that across participants, a higher percentage of frontal electrodes were classified as nested oscillations compared to those in the temporal lobe (*t*(53) = 5.55, *p* < .001, two-sample t-test of percent nested oscillations in each region), and the same relationship was demonstrated for electrodes in the parietal lobe compared to those in the temporal lobe (*t*(44) = 2.42, *p* < .05; Figure 5b).

**Figure 5.**
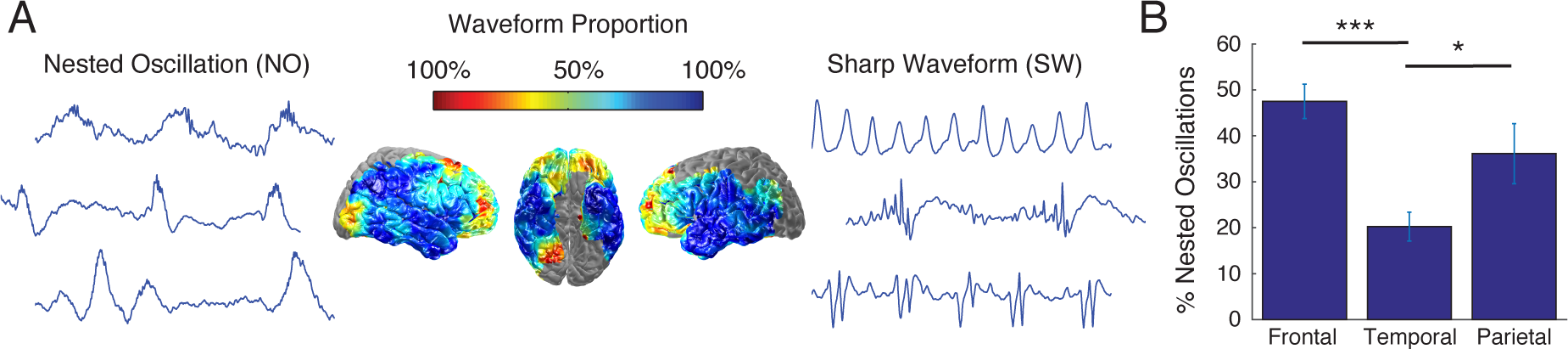
Waveform classification (**A**) Waveform distribution across brain regions. Colors correspond to percentage of electrodes demonstrating each waveform category, where warmer and cooler colors represent more NOs and SWs respectively. Exemplar iEEG traces are shown for NO (*left*) and SW (*right*). (**B**) Bar plot comparing frontal, temporal, and parietal lobes for % NO. Error bars here represent the SEM across all participants who had electrodes in those lobes. (***) indicates *p* < .001 and (*) indicates *p* < .05.

Our data suggest that sharp waveforms are generally composed of highly stereotyped and cyclostationary signals, whereas nested oscillations are more rapidly fluctuating and volatile. To quantify these differences, we examined sample entropy, a measure of signal complexity, of the unfiltered iEEG traces. Such entropy analyses have been successfully used for discerning differences in EEG signal qualities for Alzheimer’s disease and autism spectrum disorder (Mizuno et al., 2010; Catarino, Churches, Baron-Cohen, Andrade, & Ring, 2011), and thus we hypothesized that nested oscillations and sharp waveforms would be separable based on their intrinsically different waveform characteristics. Lower values of sample entropy are expected in predictable signals that contain frequent repetition of self-similar data sequences, as with the repeated low frequency components we observed in sharp waveforms. Consequently we expected that nested oscillations would exhibit higher sample entropy than sharp waveforms. We calculated the sample entropy for all trials (see Methods), and found a marked similarity between the distribution of nested oscillations and sample entropy across the cortical surface (Figure 6a). When we compared the frontal, temporal, and parietal lobes, we found that the frontal and parietal lobes had significantly higher sample entropy than the temporal lobes (frontal to temporal *t*(59) = 3.54, *p* < 0.001 and parietal to temporal *t*(54) = 3.40, *p* < 0.01; Figure 6b), tracking the observed anatomic distribution of the waveform categories (Figure 5b). Accordingly, we found that the sample entropy of nested oscillation electrodes was significantly higher than for sharp waveform electrodes across all participants (*t*(32) = 8.22, *p* < .001), suggesting that the broad categories identified by our waveform classification algorithm indeed had different salient features.

**Figure 6.**
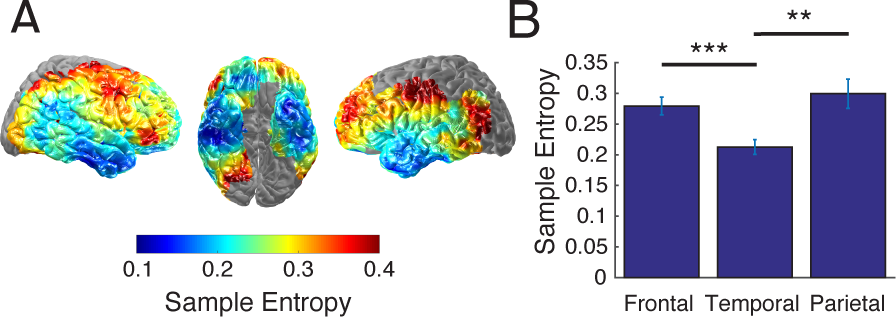
Entropy characterization (**A**) Entropy distribution across brain regions. Colors correspond to the magnitude of average sample entropy, where cooler and warmer colors represent lower and higher values respectively. (**B**) Bar plots comparing frontal, temporal, and parietal lobes for sample entropy. Error bars here represent the SEM across all participants who had electrodes in those lobes. (***) indicates *p* < 0.001 and (**) indicates *p* < 0.01.

Given the anatomic distribution of nested oscillations and sharp waveforms, we then asked how memory related changes in measured PAC in these two electrode categories compared to the changes observed with theta to low and high gamma PAC (Figure 1d). When only considering electrodes demonstrating nested oscillations, we found a significant decrease in theta to high gamma PAC during successful encoding in the left inferior frontal gyrus (*p* < .05, permutation procedure, corrected for multiple comparisons; Figure 7a). This anatomic region overlapped with the left frontal change in theta to high gamma PAC identified when we examined memory related changes irrespective of waveform type. These data therefore suggest that the memory effects observed in theta to high gamma PAC are largely carried by changes in nested oscillations that only occur in approximately half of the high gamma electrodes. Conversely, when considering only electrodes demonstrating sharp waveforms, we observed significant decreases in theta to low gamma coupling in the left temporal lobes during both encoding and retrieval (*p* < .05, permutation procedure, corrected for multiple comparisons; Figure 7b). This pattern overlapped with the observed changes in theta to low gamma PAC seen independent of waveform type, and is consistent with our finding that the vast majority of changes in theta to low gamma PAC are mediated by changes in sharp waveforms. Interestingly, for theta to high gamma PAC, it appears that the significant right hemispheric cluster identified in Figure 1d has contributions from both sharp waveforms in the superior temporal gyrus and nested oscillations in the inferior frontal gyrus, although neither of these areas are significant alone (Figure 7).

**Figure 7.**
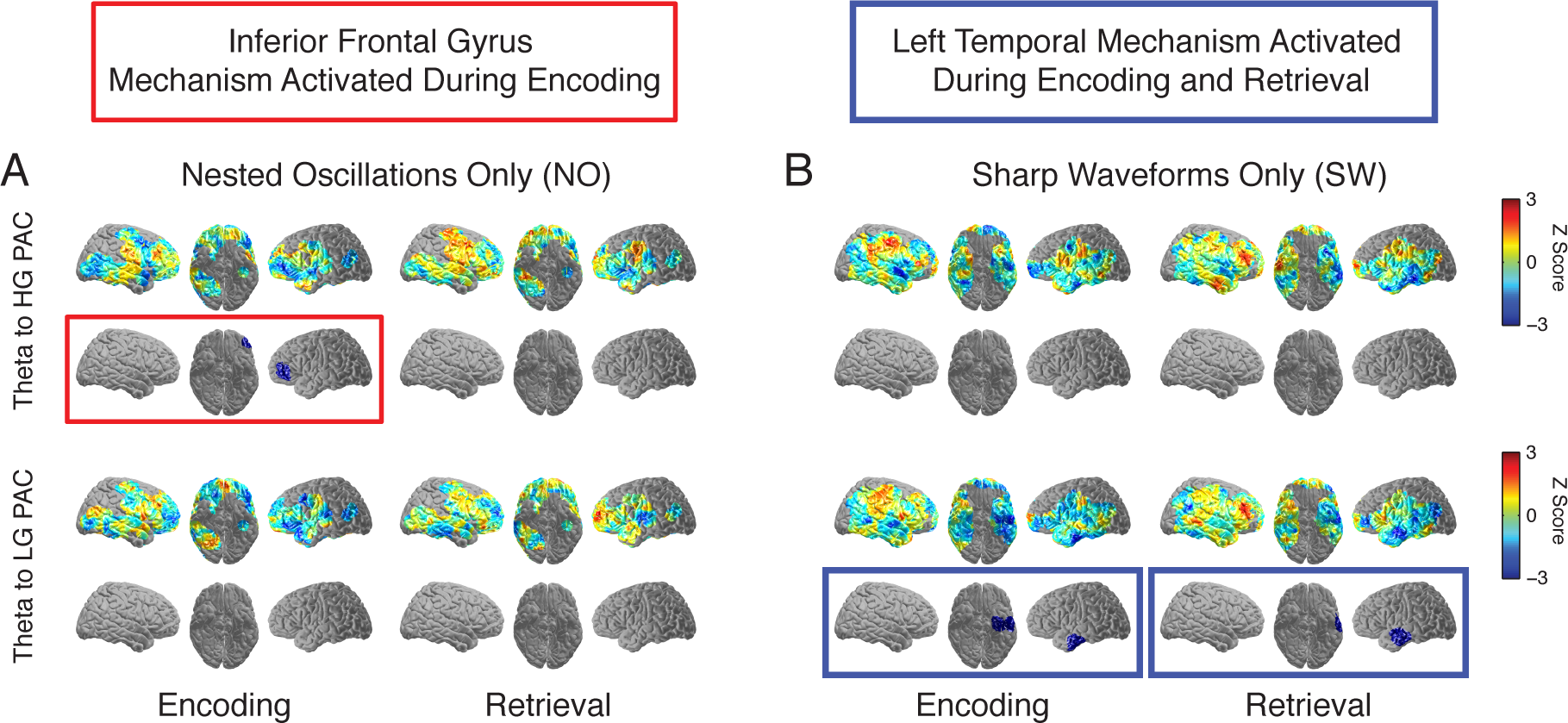
Whole brain analysis across participants for theta band PAC for correct versus incorrect trials. (**A**) Theta to HG PAC. (**B**) Theta to LG PAC. Colors in the first and third rows correspond to the average *z*-score differences between correct and incorrect trials in each ROI during encoding and retrieval. ROIs exhibiting significant differences, corrected for multiple comparisons, are shown in the second and fourth rows. The blue color of the significant regions in these rows indicates PAC was significantly less for correct trials compared to incorrect trials.

## Discussion

Our data demonstrate that successful memory encoding and retrieval are associated with significant decreases in a measure of cortical phase-amplitude coupling (PAC), and these decreases were anatomically and functionally segregated among several memory relevant areas. Using a waveform analysis to investigate the electrophysiological contributions to PAC, we found two waveform morphologies, nested oscillations and sharp waveforms, that were also anatomically segregated across the cortex and tracked the observed changes in apparent PAC. Our data suggest that these different waveform morphologies may therefore represent different underlying neuronal mechanisms that each contribute to the observed decreases in measured cortical PAC during human episodic memory formation.

While most interpretations of PAC have suggested that increases are functionally significant (J. Lisman, 2005; Canolty et al., 2006; Tort et al., 2008; Canolty & Knight, 2010), recent studies have also demonstrated the role of decreases in PAC for successful memory formation in the human hippocampus (Lega et al., 2014; Axmacher et al., 2010; Leszczynski et al., 2015). Our finding that brain regions demonstrate significant decreases in PAC as a function of episodic memory (Figure 1) are therefore among competing claims about the functional role of PAC in human cortex. One possibility is that neuronal over-coupling, manifest as PAC, may serve to inhibit, rather than enhance, effective neural processing (Hanslmayr, Staresina, & Bowman, 2016).

Over-coupling has been identified in pathologic conditions such as Parkinson’s disease (de Hemptinne et al., 2013; Yang, Vanegas, Lungu, & Zaghloul, 2014), and therapeutic deep brain stimulation that disrupts this over-coupled state has been shown to improve clinical symptoms (de Hemptinne et al., 2015). From a conceptual framework, these over-coupled, stereotyped neuronal phenomena have been described as informational lesions (Grill, A., & Miocinovic, 2004; Voytek & Knight, 2015). Hence, the reductions in coupling that we observe suggest that phase locked cortical activation may actually be poor for memory encoding and retrieval (Hanslmayr et al., 2016). While hippocampal PAC may be important for organizing memory representations (J. Lisman, 2005; Axmacher et al., 2010; Leszczynski et al., 2015), such coupling may limit flexibility in cortical neural processing, and therefore in information processing capabilities.

Interpreting these changes in PAC is difficult, however, as the precise neurophysiological contributions to measured PAC in the context of memory encoding and retrieval are complex (Kramer et al., 2008; Aru et al., 2015; Jones, 2016). Our data are consistent with previous reports that suggest measures of human cortical PAC can be derived from repeated nested oscillations (Canolty et al., 2006; He et al., 2010) or from repeated non-sinusoidal signals (Kramer et al., 2008; Aru et al., 2015; Jones, 2016; Lozano-Soldevilla et al., 2016). Indeed, the extent of measured PAC, both in our recorded iEEG signals and in simulated and non-neural traces (Tort et al., 2008, 2013; He et al., 2010; Aru et al., 2015), is related to the strength and consistency of characteristic and repeated waveforms in the time series trace. The constituent frequency components of the underlying stereotypical waveform determine the modulating and modulated frequencies observed in measures of PAC (Aru et al., 2015; Burke et al., 2015; Jones, 2016). And the preferred phase is related to the temporal arrangement of these functional components.

In this sense then, the functional significance of measures of PAC during memory encoding and retrieval may be related to the underlying iEEG waveforms themselves. We examined differences in the strength and consistency of these waveforms and found that correct encoding and retrieval caused significantly reduced phase-triggered average waveform amplitudes in the left temporal lobe, while correct encoding alone caused reduced average waveform amplitudes in the left frontal lobe (Figure 3). These observed differences in amplitude were similar, although not identical, in extent and anatomical location to the effects measured with PAC, and raise the possibility that these changes in the iEEG signal may represent a neural correlate of episodic memory. While measures of PAC may be difficult to interpret, their utility could be in identifying the presence of these underlying waveforms, and the neural mechanisms they may represent.

As such, we used an automated classification algorithm to gain insight into the spatial distribution of the constituent components that contribute to the observed measures of PAC. Nested oscillations were more prevalent in the frontal lobes than in the temporal lobes. Conversely, measures of PAC in the temporal lobe, as well as those associated with alpha oscillations, appear to have a stronger link with sharp waveforms that do not demonstrate the presence of a clear high gamma oscillation (Figure 5). It is possible, of course, that our automated approach is simply incapable of detecting all nested gamma oscillations, and the distinction between true nested oscillations and neural stochastic volatility remains an area of active investigation (Lachaux, Axmacher, Mormann, Halgren, & Crone, 2012; Burke et al., 2015; Aru et al., 2015; Jones, 2016). However, we examined signal complexity and found that it, too, had an anatomic distribution that was closely related to the distribution of waveform types (Figure 6). Together, these data suggest that the waveform characteristics of the frontal lobes are different than those of the temporal lobes, and this in turn provides a basis for interpreting the memory related modulation of activity in these areas.

Indeed, when we examined the memory related changes in electrodes classified as either containing nested oscillations or sharp waveforms, we found that they too were anatomically segregated, suggesting the existence of different neuronal mechanisms underlying successful memory formation (Figure 7). Nested oscillations exhibited significant decreases during successful encoding that were confined to only the left inferior frontal gyrus and the theta to high gamma frequency bands. Hence, our data suggest that nested oscillations are the primary contributor to the memory related changes we observe in higher frequency PAC. We found that electrodes characterized by sharp waveforms, on the other hand, exhibited significant decreases in theta to low gamma coupling in the left temporal lobe. Correct retrieval involved the same left temporal lobe regions, suggesting that the observed memory related changes in low gamma PAC are largely driven by sharp waveforms.

The anatomic and functional segregation of both waveform types and frequency bands involved in the observed measures of PAC suggest the presence of different underlying neuronal mechanisms in different anatomic regions that contribute to human episodic memory formation. Previous studies suggest nested oscillations may play an important role in executive control and working memory maintenance in the prefrontal cortex (Voytek et al., 2015). The reduction in PAC for nested oscillations, however, may be due to extended duty cycles of high frequency oscillations (Jensen, Gips, Bergmann, & Bonnefond, 2014). In this framework, PAC decreases primarily because gamma cycles are distributed over more phase bins of the modulating frequency. Similar changes have been observed in monkey visual cortex (Spaak, Bonnefond, Maier, Leopold, & Jensen, 2012) and in human hippocampus (Leszczynski et al., 2015; Heusser et al., 2016), and may underlie the reductions in PAC we observe in the left inferior frontal gyrus during encoding.

Conversely, sharp deflections in the iEEG waveform result in increases in broadband power that have been shown to reflect neural activation (Makeig et al., 2002; Manning et al., 2009; Miller et al., 2007; Burke et al., 2015). Phase-locked repetition of these deflections may therefore entrain neural activity to a single rhythm in the temporal lobes. In this case then, successful memory formation may require a reduction in the extent to which neural activation is entrained to a low frequency signal, allowing greater flexibility in neural processing. Our data demonstrating decreases in the phase-triggered average waveform amplitude support this possibility.

Together, our data demonstrate task-related decreases in the extent of both nested oscillations and sharp waveforms that correlate with successful memory encoding and retrieval. Using a systematic method to distinguish these waveforms, we show that memory related changes in measures of cortical PAC may be related to distinct underlying neural mechanisms represented by these different waveforms. Hence, although the proper interpretation of PAC and its functional significance still remain unclear (Kramer et al., 2008; Canolty & Knight, 2010; Lachaux et al., 2012; Burke et al., 2015; Aru et al., 2015; Jones, 2016), measures of PAC may be important in informing us about the complex nature of underlying waveforms in the iEEG signal and their changes during successful memory encoding and retrieval.

## Author Contributions

A.P.V. and K.A.Z. designed research; A.P.V., R.B.Y., J.H.W., S.K.I., and K.A.Z. performed research; A.P.V. analyzed data; A.P.V., R.B.Y., J.H.W., S.K.I., and K.A.Z. wrote the paper.

## Acknowledgements

We thank John Burke, Julio Chapeton, Baltazar Zavala, John Cocjin, and Rafi Haque for helpful and insightful comments on the manuscript. This work was supported by the Intramural Research Program of the National Institute for Neurological Disorders and Stroke. This work was also supported in part by the National Institute of General Medical Sciences (NIGMS) grant T32 GM007171 to APV. We are indebted to all patients who have selflessly volunteered their time to participate in this study.

The authors declare no competing financial interests.

